# Hospital outbreak of carbapenem-resistant Enterobacteriales associated with an OXA-48 plasmid carried mostly by *Escherichia coli* ST399

**DOI:** 10.1101/2020.06.15.148189

**Authors:** Alice Ledda, Martina Cummins, Liam P. Shaw, Elita Jauneikaite, Kevin Cole, Florent Lasalle, Deborah Barry, Caryn Rosmarin, Sudy Anaraki, David Wareham, Nicole Stoesser, John Paul, Rohini Manuel, Benny P Cherian, Xavier Didelot

## Abstract

A hospital outbreak of carbapenem-resistant Enterobacteriales was detected by routine surveillance. Whole genome sequencing and subsequent analysis revealed a conserved promiscuous OXA-48 carrying plasmid as the defining factor within this outbreak. Four different species of Enterobacteriales were involved in the outbreak. *Escherichia coli* ST399 accounted for 20/55 of all the isolates. Comparative genomics with publicly available *E. coli* ST399 sequence data showed that the outbreak isolates formed a unique clade. The OXA-48 plasmid identified in the outbreak differed from other known plasmids by an estimated five homologous recombination events. We estimated a lower bound to the plasmid conjugation rate to be 0.23 conjugation events per lineage per year. Our analysis suggests co-evolution between the plasmid and its main bacterial host to be a key driver of the outbreak. This is the first study to report carbapenem-resistant *E. coli* ST399 carrying OXA48 as the main cause of a plasmid-borne outbreak within a hospital setting. This study supports complementary roles for both plasmid conjugation and clonal expansion in the emergence of this outbreak.

## Introduction

Horizontal Gene Transfer (HGT) is a key feature of bacterial adaptation (Frazão *et al.*, 2019). HGT of genes from neighbouring organisms helps bacteria to adjust to environmental change. Acquisition of antimicrobial resistance (AMR) through HGT represents a serious threat to public health (MacLean and San Millan, 2019). HGT allows the spread of AMR genes within a single generation of bacteria. Bacterial host sensitivity for HGT may be narrow, with transfer limited to a single species, or broad, with spread of resistance genes between multiple species. Horizontal transfer of gene clusters may propagate multidrug-resistant (MDR) strains.

Plasmids are vectors of HGT, found ubiquitously in bacterial species (Norman, Hansen and Sørensen, 2009; Mathers *et al.*, 2015). Conjugative plasmids are able to transfer copies of themselves to neighbouring bacteria and have been described as carriers of specific AMR genes (Lanza *et al.*, 2014; Mathers *et al.*, 2015; Sheppard *et al.*, 2016). One example is carbapenemases: genes which can confer resistance to carbapenems, a class of beta-lactam antibiotics(León-Sampedro *et al.*, no date; Poirel, Potron and Nordmann, 2012). These antibiotics are widely used in clinical practice as they are active against extended-spectrum β-lactamase ESBL-producing Enterobacteriaceae which are resistant to other beta-lactams (Potron *et al.*, 2011; Nordmann, Dortet and Poirel, 2012). Acquisition by bacteria that also produce ESBL may render them resistant to all beta-lactams currently used for treatment (Nordmann, Dortet and Poirel, 2012; Poirel, Potron and Nordmann, 2012; Skalova *et al.*, 2017). Several classes of carbapenemase have been described, and genes are typically found on plasmids and other mobile genetic elements (MGEs) (Potron *et al.*, 2011; Nordmann, Dortet and Poirel, 2012; Poirel, Potron and Nordmann, 2012; Findlay, Hopkins, Alvarez-Buylla *et al.*, 2017; Skalova *et al.*, 2017).

pOXA-48 is a typical example of a plasmid carrying a gene (*bla*_OXA-48_) conferring resistance to carbapenems (Poirel, Potron and Nordmann, 2012; Skalova *et al.*, 2017). pOXA-48 has been found in a wide range of Enterobacteriales (León-Sampedro *et al.*, no date; Mahon *et al.*, 2019). This presents a public health challenge: once acquired, the plasmid can spread among the Enterobacteriales present in the gut microbiome, including potential pathogens. In this paper, we present a genomic analysis of an outbreak of carbapenem-resistant Enterobacteriales driven by pOXA-48 in a hospital ward in the United Kingdom. Previous studies have reported multi-species outbreaks of pOXA-48 in hospital settings (David *et al.*, no date; León-Sampedro *et al.*, no date; Pulss *et al.*, 2018; Hernández-García *et al.*, 2019), but have remained largely descriptive. Here, we introduce a mathematical model of plasmid transmission between different host STs and species to obtain a lower bound for the number of conjugation events per year.

## Results

### Outbreak overview

A total of 55 carbapenem-resistant isolates were cultured from 48 unique patients from the first 10 months of the outbreak (May 2016 to February 2017). From both *in vitro* assays and subsequent whole genome sequencing, all carried the OXA-48 gene on a pOXA-48 plasmid (see below). Four different species of Enterobacteriales were identified (Figure 1). Most isolates were *Escherichia coli* (n=43), then *Klebsiella pneumoniae* (n=7), *Enterobacter cloacae* (n=4) and *Citrobacter koseri* (n=1).

**Figure 1.**
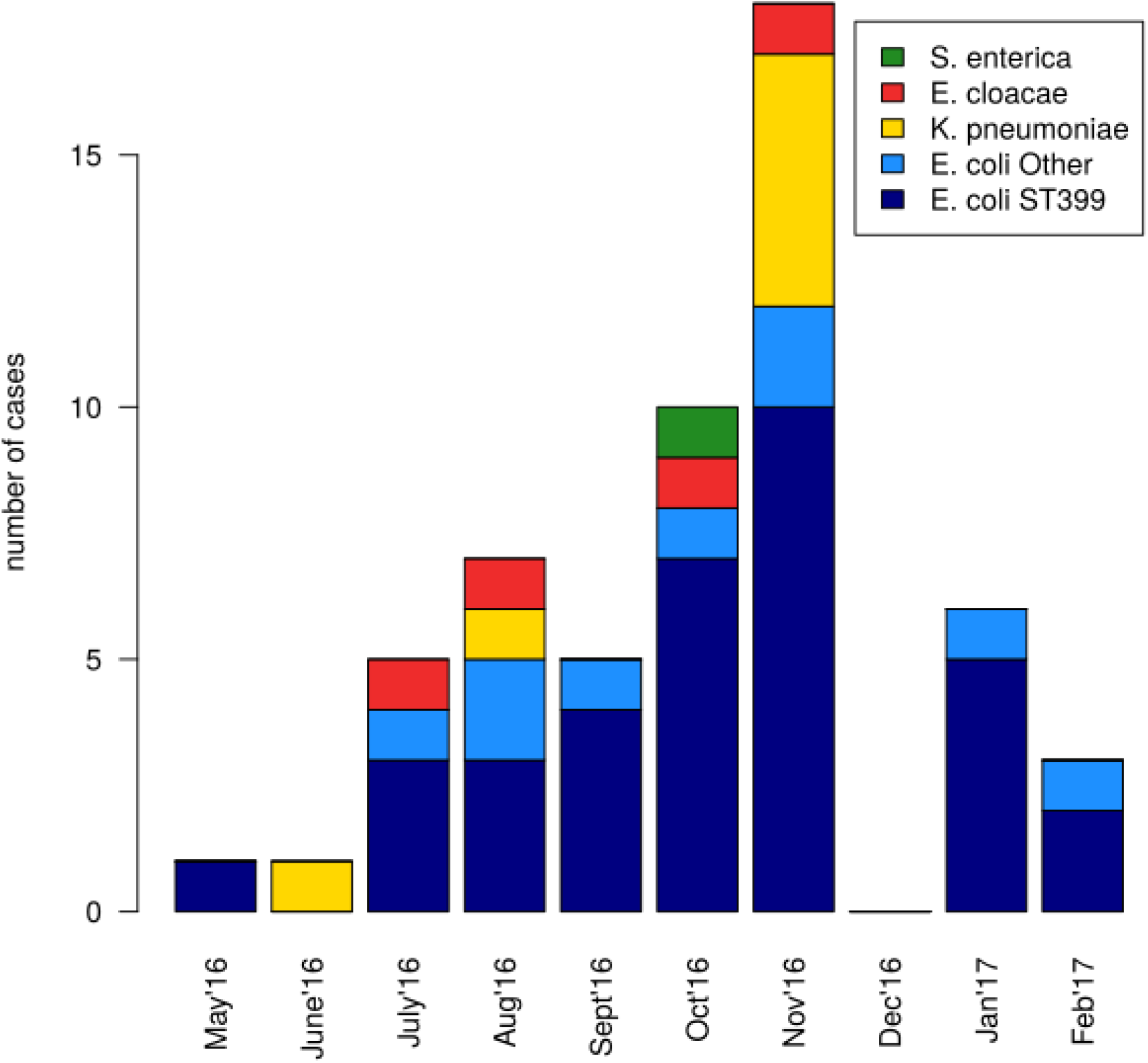
Progression of the OXA-48 outbreak, the main bacterial isolates and MLST types involved are highlighted in different colors.

35 of the 43 *E. coli* isolates (81%) belonged to ST399 according to the Achtman multi-locus sequence typing (MLST) scheme of *E. coli* (Clermont *et al.*, 2013; Dale and Woodford, 2015). Eight other *E. coli* strains belonged to different STs, including two novel STs (Table Supplementary1). *E. coli* ST399 has previously been described as a carrier of the pOXA-48 plasmid (Findlay, Hopkins, Alvarez-Buylla *et al.* 2017), although it was not a main host, only 2 instances of carriage were reported. None of the OXA-48-positive *E. coli* isolates were ST38, which was previously found associated with pOXA-48 (Turton *et al.*, 2016). Three of the seven *K. pneumoniae* isolates were novel STs and none of them were ST14 or ST10, previously found associated with pOXA-48 carriage (Findlay, Hopkins, Loy *et al.* 2017). Only one of the four *E. cloacae* isolates belonged to a previously described ST.

The outbreak index case carried an *E. coli* isolate belonging to ST399 (Figure 1) sampled at the end of May 2016. The second case yielded an *K. pneumoniae* isolate sampled in mid June 2016. Due to the increased number of patient cases affected by OXA-48-positive Enterobacteriaceae, the ward was closed to new patient admissions in December (hence the lack of cases during this month in figure 1). The ward was re-opened after the Christmas holidays. Thereafter, only *E. coli* cases were detected.

### Highly conserved pOXA-48 plasmid

Plasmid pOXA-48 is a 61,881bp plasmid belonging to the L/M incompatibility group (IncL/M), carrying only replication machinery and the OXA-48 carbapenem resistance determinant (Bonnin *et al.*, 2013). All of the 55 samples were confirmed in-silico to be carrying the OXA-48 allele, as previously described (no OXA-181 or OXA-163, which have a similar phenotype, were found) (Poirel *et al.*, 2011; Potron *et al.*, 2011). All the OXA48 genes were found on a p-OXA48-like plasmid. No insertions or deletions with respect to the previously described pOXA-48 reference (NCBI GenBank accession JN626286) (Poirel, Bonnin and Nordmann, 2012) were found by any of the approaches used. The genomic content of the pOXA-48 plasmid was highly conserved.

The alignment of all pOXA-48 plasmids identified in the study was used to build a phylogenetic tree in which the reference pOXA-48 was used as an outgroup (Figure 2a). Bacterial species are intermixed within the phylogeny tree (Figure 2a), indicating that the outbreak is not caused by the separate clonal spread of each bacterial species.

**Figure 2.**
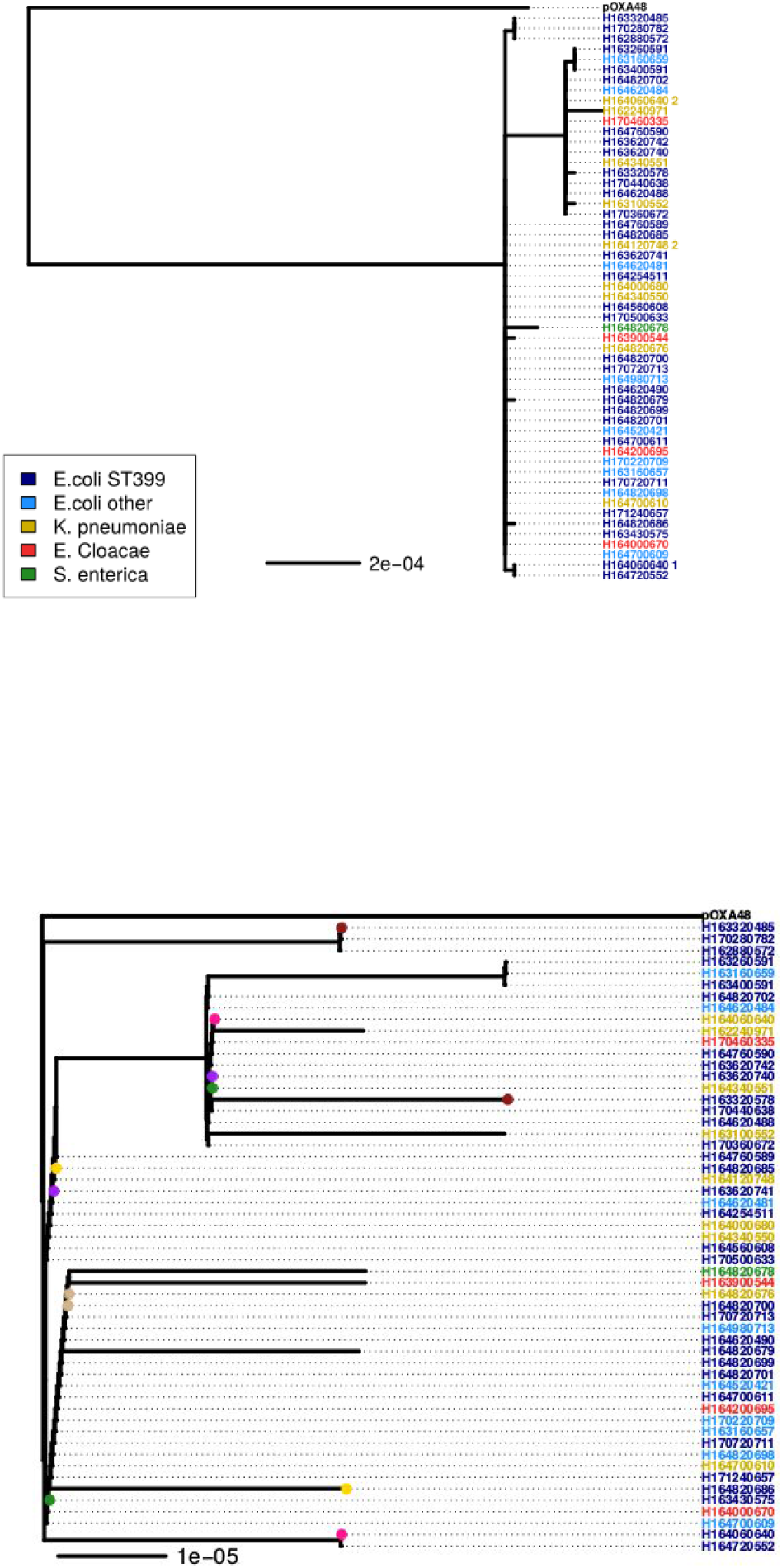
Phylogenetic analysis of the pOXA-48 plasmids. A. Phylogenetic tree of all the pOXA-48 sequences estimated with PhyML under the GTR model, with tip colors representing bacterial species. B. Clonal genealogy estimated by ClonalFrameML.

### Recombination analysis

ClonalFrameML was used to identify the recombination events among the studied plasmids (Figure 2b). The overall tree topology remained unchanged once recombined segments of the plasmid were removed. The reference pOXA-48 plasmid remains genetically separate from all other plasmids in the tree (Figure 2b).

To understand the evolutionary history of the plasmids identified during the outbreak, all nucleotide differences between any pair of plasmids were identified (including the reference pOXA-48). The mutations and their abundance are shown in Figure 3. A total of 155 nucleotide variants were found, 129 of which distinguished the pOXA-48 reference from all other plasmids in the dataset (Figure 3 b,c).

**Figure 3.**
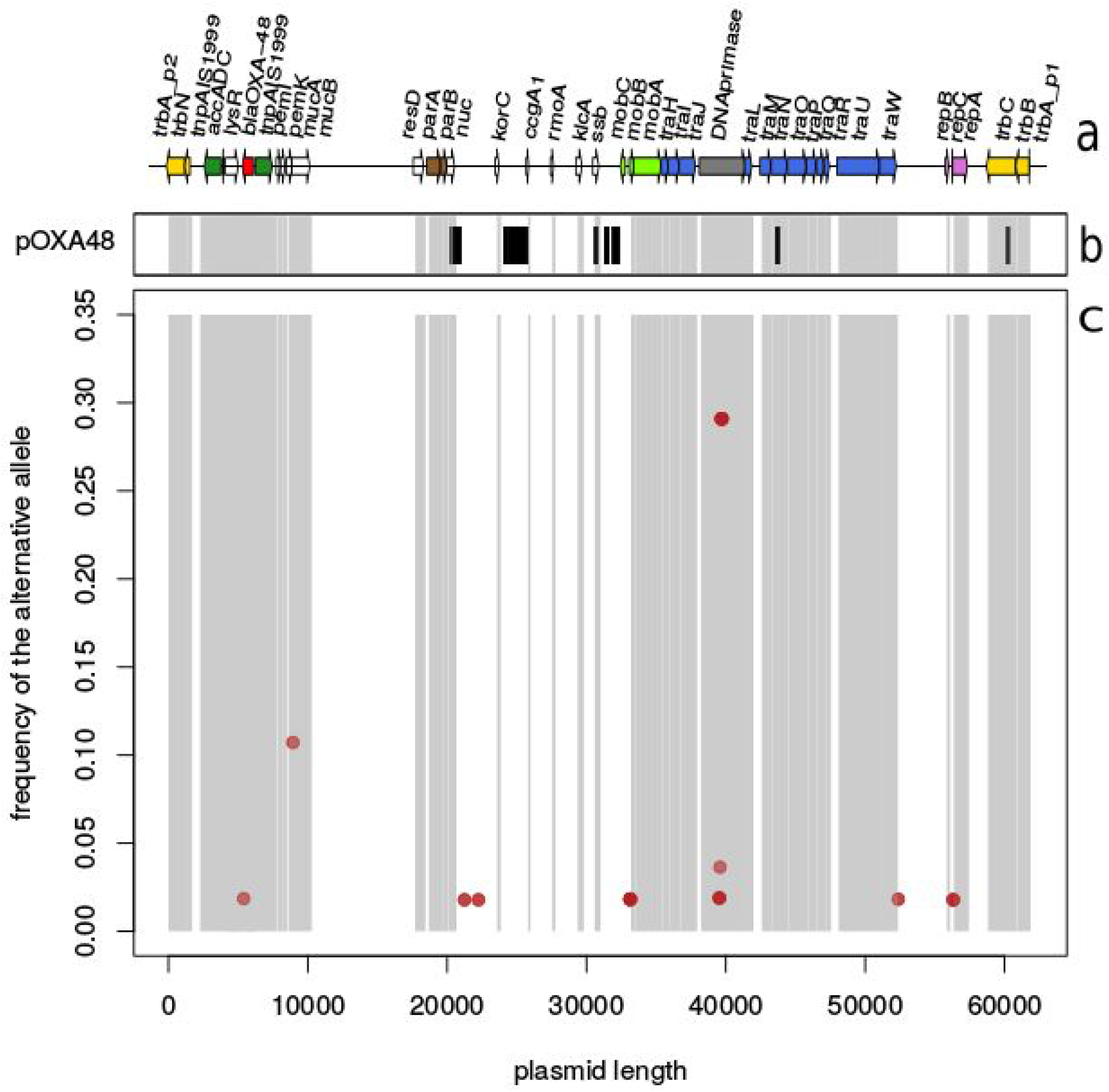
Genomic distribution of polymorphisms. Panel (a) shows the plasmid genetic content. Panel (b) shows the mutations distinguishing the reference plasmid pOXA-48 from the rest of the plasmids in this study. Panel (c) shows the mutations present only in plasmids included in this study and their abundance (y-axis). The shadowed areas in panels (b) and (c) correspond to coding regions in the plasmid (panel (a)).

In non-recombining sequences with weak mutational biases, neutral mutations are expected to be distributed almost uniformly along the sequence. This is clearly not the case of the variants which distinguished outbreak plasmid sequences from the pOXA-48 reference, which were localised in five clusters (Figure 3b), suggesting several independent homologous recombination events occurred. Four out of five of these putative events are located in coding regions, and one (the second from the left) is located in an intergenic region (Table 1). Most of the variants lie in intergenic regions (114 out of 129), with the majority concentrated in the second recombinant locus. Of the 15 mutations located in coding regions, 10 are synonymous and 5 are non-synonymous.

**Table 1:** Nucleotide differences between the sampled plasmids and pOXA-48, see also upper panel of Figure 3.

| recombinant<br>region (bp) | number of<br>SNPs | gene |  |  |  |
| --- | --- | --- | --- | --- | --- |
|  |  |  | intergenic | syn | non-syn |
| 20514 - 20961 | 11 | *nuc* | 5 | 4 | T31A, G174D |
| 24206 -25674 | 95 | - | 95 | - | - |
| 30661 - 32310 | 17 | *ssb* | 14 | 2 | R27G |
| 43636 - 43837 | 4 | *traN* | - | 2 | Q89R, T156G |
| 60324 - 61717 | 2 | *trbC* | - | 2 | - |

Overall, the plasmid gene content is highly conserved (Supplementary figure 1). No single nucleotide variant is observed outside of the putative recombination events, suggesting no nucleotide substitution occurred during the diversification of the plasmids in our sample. Considering the length of the plasmid and the time elapsed since the sampling of the reference pOXA-48 plasmid we would have expected to see very few, if any, nucleotide substitutions (see Methods section “Expected number of mutations”). Similarly, we observed no structural variants such as insertion/deletion of insertion sequences, transposon or any other genetic material

### Outbreak dynamics and estimation of inter-host conjugation rate

The predominant bacterial host *E. coli* strain ST399 was isolated repeatedly throughout the outbreak.

The observed distribution of hosts for pOXA-48 suggests a parsimonious scenario where an ongoing clonal epidemic of *E. coli* ST399 carrying the pOXA-48 plasmid, with multiple independent conjugation events of pOXA-48 into other hosts (both other *E. coli* STs and other species). To test whether this description was consistent with predictions of a population genetics theoretical framework, we built a Markov model of plasmid transmission between hosts, assuming one founder host (see Methods section “Neutral model of inter-bacterial-host conjugation”).

The Markov model confirms that the founder host of this outbreak was *E. coli* ST399 (se Methods). The model predicts that plasmids found in *E. coli* ST399 have been either vertically inherited or obtained by intra-host conjugation from other ST399 *E. coli* strains, as conjugation from another host into *E. coli* ST399 was highly unlikely (see Methods section “Expected number of mutations”). This model was then used to infer the underlying evolutionary scenario following a host-switching process under an “infinite-hosts” conjugation model.

We used this mathematical model of conjugation to draw conclusions on the evolutionary history of the outbreak based on the host frequency spectrum of pOXA-48 (Ferretti *et al.*, 2017). The host frequency spectrum is composed of singletons, except for *E. coli* ST399, which was also both the first identified and thus considered to be the ancestral host. Two evolutionary scenarios are consistent with this pattern: i) *E. coli* ST399, in combination with pOXA-48, had an evolutionary advantage compared to other host-plasmid combinations (‘co-evolution’ scenario), or ii) their association resulted from the recent rapid expansion of *E. coli* ST399 (‘neutral association’ scenario). The hypothesis of co-evolution of pOXA-48 with ST399 implies that the plasmid has reduced fitness in other bacterial hosts. It follows that the observation of the plasmid in all the other bacterial hosts, which appear as singletons in the host frequency spectrum, were most likely derived from very recent conjugation events and have not yet been purged by purifying selection (Wen-Hsiung, 1993; Ferretti *et al.*, 2017). In the ‘neutral association’ scenario, the rapid expansion of the plasmid population would generate a star-like genealogical tree for the plasmids. In this scenario each different plasmid lineage evolved and conjugated independently after the start of the outbreak (Ferretti *et al.*, 2017). The inter-host conjugation rate under the assumption of the neutral association scenario is easily inferred. In this scenario, the plasmid tree would be approximately star-like and all non-ST399 *E. coli* hosts result from independent conjugation events. All conjugation events would have occurred at random times between the present and the time of the most recent common ancestor (tMRCA) (see Methods section “Estimation of pOXA-48 conjugation rate”). In our dataset, BactDating (Didelot *et al.*, 2018) estimates that the most recent common ancestor of the *E. coli* ST399 tree is around late 2013 (supplementary figure 1). Using this information, we estimate a pOXA-48 conjugation rate of about 0.23 conjugation events per lineage per year.

### A case of co-evolution between plasmid and host?

The genomic data available could be explained by both the recent neutral expansion scenario or the co-evolution scenario or any scenario in between. Although we used the neutral scenario hypothesis to estimate a lower bound to the conjugation rate in our sample, the co-evolution hypothesis cannot be discarded. The prospect of co-evolution between *E. coli* ST399 and the plasmid responsible for the outbreak was therefore investigated in more detail. To provide context to our observations, we checked whether the studied outbreak might be part of a broader epidemic of *E. coli* ST399. We searched EnteroBase to obtain all publicly available *E. coli* ST399 genomes and found a total of 82 genomes. These genomes were mapped to the *E. coli* ST399 reference genome and to the reference plasmid pOXA-48 (see Methods).

The search for the plasmid in the downloaded dataset yielded inconclusive results as no single full-length pOXA-48 plasmid was found.No OXA-48 gene was found to occur within the publicly available genomes of *E. coli* ST399 (n=79); only partial fragments of the plasmid that do not include the resistance gene OXA48 were found in 20 of these genomes.

A phylogenetic tree of *E. coli* ST399 including the reference, the samples included in the present study and the genomes from EnteroBase is shown in Figure 4. It is obtained using whole genome, on reference mapping of the raw data downloaded from EnteroBase. All the samples from this study except two are clustered in a single clade, pointing to an expanding outbreak of vertically inherited pOXA-48. The two samples not clustering in the main clade were sequenced at the end of the sampling period: in January and February 2017. None of the other EnteroBase sequences formed clusters within the same clade, indicating the outbreak had a single main source associated to the hospital ward reported in this study.

**Figure 4.**
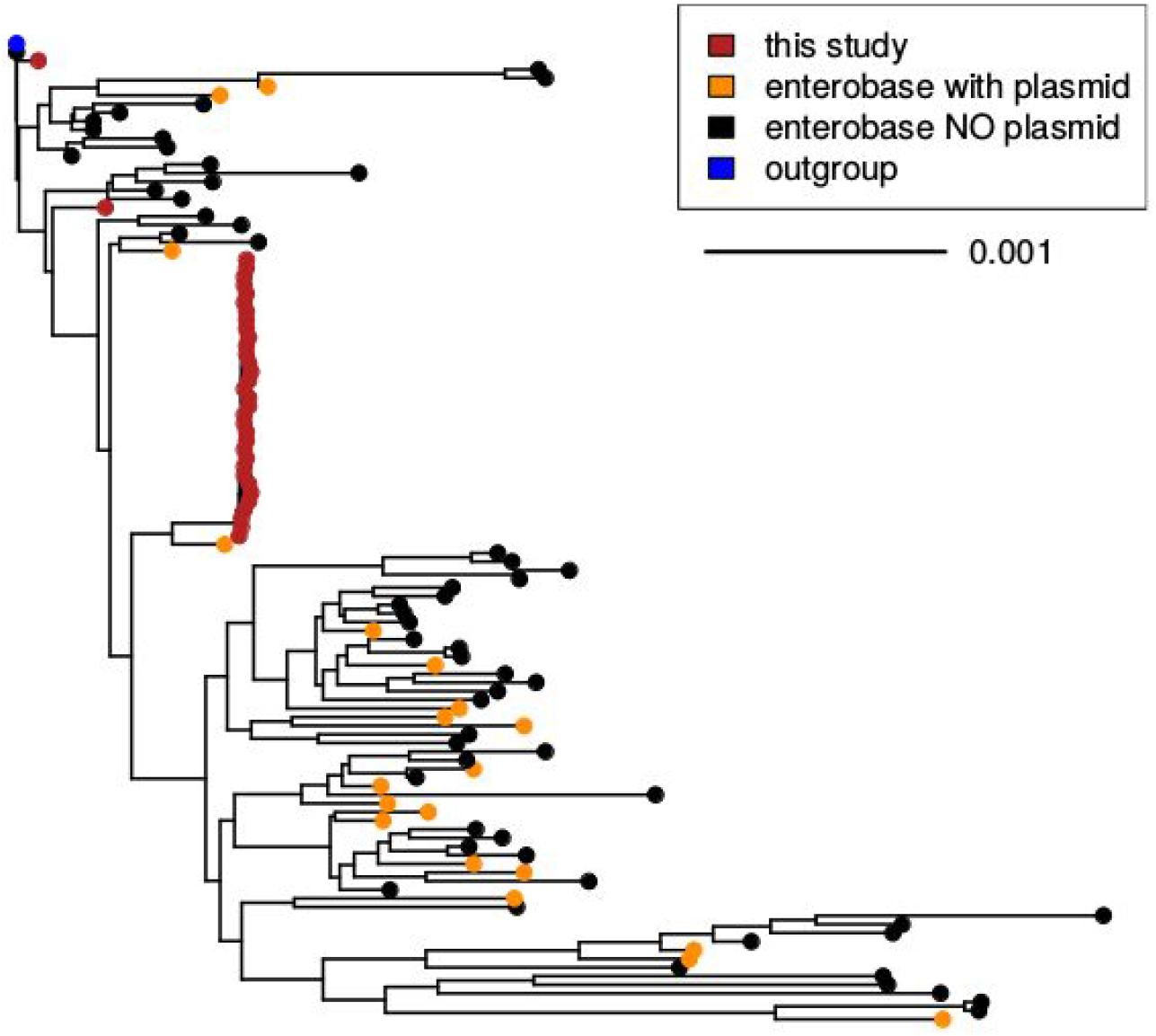
Phylogenetic tree of *E. coli* ST399. In orange and black the samples from EnteroBase, in orange the ones in which we found traces of the plasmids (although no complete plasmid was found in any of the samples) and in black the samples in which we did not find any trace of the plasmid, in blue the reference genome and in red the samples included in this study. Note that all except two of the samples from this study cluster exclusively into a single clade pointing to a clonal expansion of this specific clade.

## Discussion

Genomic surveillance of AMR pathogens is of paramount importance, as it allows the early identification of plasmid-borne AMR genes transmission, and thus to prevent further dissemination. This systematic approach is particularly crucial to prevent plasmid transmission to bacterial hosts that are already resistant to certain antibiotics, which would result in the creation of multi-resistant strains and to potentially untreatable infections. In this paper we present the analysis of an outbreak caused by plasmid, carrying antimicrobial resistance genes, detected by surveillance in a hospital ward. The outbreak involves different Enterobacteriales species and covers the first 10 months of the 18-months long outbreak. Analysis of sampled sequences suggested a clear outbreak dynamic: there is a clonal expansion of pOXA-48 carried by *E. coli* ST399, while simultaneously the plasmid conjugates into *E. coli* belonging to other STs as well as other Enterobacteriales species. This description is supported by a population genetics model based on the frequency spectrum of bacterial hosts. Such outbreak dynamics are typical of pOXA-48, although the main host is usually from the genus *Klebsiella* (Findlay, Hopkins, Alvarez-Buylla *et al.* 2017; Findlay, Hopkins, Loy *et al.* 2017). The plasmids were highly conserved, in accordance to previous pOXA-48 outbreaks (León-Sampedro *et al.*, no date; Skalova *et al.*, 2017; Pulss *et al.*, 2018; Hernández-García *et al.*, 2019; Mahon *et al.*, 2019). We were able to assemble the full-length sequence of the plasmid using Illumina short-read sequencing, without the need for long read sequencing.

Contrary to previously described pOXA-48 outbreaks (León-Sampedro *et al.*, no date; Findlay, Hopkins, Alvarez-Buylla *et al.* 2017; Skalova *et al.*, 2017; Mahon *et al.*, 2019) the main host in our study was *E. coli*. More than half of the *E. coli* samples belong to *E. coli* ST399, while the others belong to different STs, including some that have not yet been characterised. *E. coli* ST38 has been described as the main host in a p-OXA48 outbreak (Turton *et al.*, 2016), but in that case the plasmid had been partially inserted in the chromosome, meaning that outbreak relied on the clonal expansion of the bacterial host. In this outbreak we found a clonal expansion of the combination of an *E. coli* ST399 genome with a pOXA-48 plasmid.

Because of the short plasmid length (~61kbp)(Poirel, Bonnin and Nordmann, 2012; Bonnin *et al.*, 2013) and relative brevity of the outbreak, not enough mutations accumulated to allow us to resolve the plasmid phylogenetic tree and ascertain the precise evolutionary scenario involved in the outbreak. It might be a ‘co-evolutionary scenario’: the association between pOXA-48 and *E. coli* ST399 is a favourable one and pOXA-48 in any other bacterial background is not fit enough and gets purged by natural selection. Or it might be a ‘neutral association scenario’: pOXA-48 underwent a recent expansion and all the conjugation events happened after the beginning of the outbreak.

In the case of a recent expansion and a star-like tree the plasmid conjugation rate is readily estimated. This rate is in fact a lower bound for the real conjugation rate in this specific outbreak. It might be a gross under-estimation of the conjugation rate if the plasmid is strongly adapted to *E. coli* ST399. In such a case, conjugation events would have occurred much more recently than estimated, by a factor given (in bacterial generations) by the inverse fitness advantage of *E. coli* ST399 compared to other hosts. The timescale of these plasmid transmission events could thus be much shorter than the time to the most recent common ancestor of all plasmids in this study, hence resulting in a much faster conjugation rate. So 0.23 per lineage per year is a lower bound to the real conjugation rate in our sample. This conjugation rate over time cannot be directly compared to *in vitro* experiments, which measure the conjugation frequency: the proportion of colonies infected over the total number of colonies during the course of an experiment. Nevertheless, such experiments confirm that pOXA-48 can be mobilized between species and genera with a high conjugation frequency (Turton *et al.*, 2016; Mahon *et al.*, 2019).

In the ‘co-evolutionary scenario’, determining the exact evolutionary scenario would not only be important to determine the exact conjugation rate, but also to infer the fitness advantage of the association of *E. coli* ST399 with pOXA-48, which is eventually a measure of how dangerous this pairing is and how strongly public health officers should react to this specific outbreak.

Surveillance combined with detailed analysis and modelling is of paramount importance for early detection and management of plasmid outbreaks. While surveillance is important in its own right, we have shown the value of whole-genome sequencing and subsequent mathematical modelling. We managed to identify the clonal part of the outbreak, which is the most important one to stop in order to end the outbreak overall. We have also estimated the conjugation rate, which is useful to infer how much spill-out to expect. Future analysis of surveillance data will be valuable, not only to detect plasmid outbreaks very early on but also to provide guides to the best way to stop those outbreaks.

## Materials and Methods

### Outbreak description

The outbreak was on a vascular ward within a new hospital build. The ward had 30 beds, consisting of 10 side rooms and five four bed bays all with en-suite facilities. In each bay the beds were a minimum of 3.6 meters from bed centre to bed centre in line with HTM. The outbreak started in May 2016, when a known patient, colonised with OXA-48 *E. coli*, was admitted into a bay within the ward. The patient was mobile and socialised with other inpatients. Our initial contact screening yielded another positive patient. An outbreak was declared in June when a number of additional cases were identified. This was the beginning of our 18 month outbreak. At the beginning of the outbreak we did not undertake routine screening for patients with carbapenem-resistant Enterobacteriaceae, however as the outbreak progressed we initiated weekly and admission screens. Over a period of 18 months we identified a total of 134 patients with a resistance linked to this outbreak, the majority being asymptomatically colonised with only three patients developing an infection. Initial standard infection and prevention control measures failed due to long durations of colonisation and frequent admissions to the ward. Stricter infection control measures were later implemented, including education of healthcare workers, extension of our screening programme using PCR testing, enhanced cleaning and emptying the ward, which all contributed to the successful control of the outbreak.

Here we analyse all the samples that were identified as carbapenem resistant during the first 10 months of the outbreak. They are 55 samples from 48 unique patients during ten consecutive months from May 2016 to February 2017. All clinical isolates showing carbapenem resistance were archived and investigated. The gene allele conferring carbapenem resistance was identified using standard *in vitro* methods as OXA-48. This was confirmed *in silico* using whole genome sequencing (WGS) data.

### Bacterial isolates and genomic sequencing

Isolates were cultured from −80°C stocks onto Columbia Blood Agar (Oxoid, Basingstoke, UK) and incubated aerobically overnight. Bacterial DNA was then extracted and sequenced according to a previously described protocol (Stoesser *et al.*, 2013). In short a combination of physical (FastPrep, MP Biomedicals, USA) and chemical (QuickGene DNA Tissue Kit S, Fujifilm, Japan) lysis was used to lyse the bacterial cells and elute DNA. The DNA concentration was measured using the PicoGreen® dsDNA Assay Kit (Waltham, Massachusetts, US). Whole-genome sequencing was performed using an Illumina HiSeq2000 that generated 100 bp paired-end reads (San Diego, California, US).

### Genomic analysis

The read assembly was done using Spades 3.11 (Bankevich *et al.*, 2012). Bacterial species identification was obtained from assembled contigs using the Nullarbor (*Website*, no date; Bankevich *et al.*, 2012) suite and confirmed using the online version of KmerFinder (Hasman *et al.*, 2014; Larsen *et al.*, 2014; Clausen, Aarestrup and Lund, 2018) available on the Center for Genomic Epidemiology website (*Center for Genomic Epidemiology*, no date a). Multilocus sequence typing (MLST) was performed using MLST on the Center for Genomic Epidemiology website (*Center for Genomic Epidemiology*, no date b).

Initial plasmid *de-novo* assembly was done using the option PlasmidSpades in Spades 3.11 (Bankevich *et al.*, 2012). Since a reference plasmid was available (GenBank accession JN626286), on-reference assembly was attempted using three different methods: lastZ (Rahmani *et al.*, 2011), mummer (Kurtz *et al.*, 2004) and samtools/bcftools (Li *et al.*, 2009; Li *et al.*, 2011). The final reference-based mapping was done using mummer (Kurtz *et al.*, 2004). Reference-based mapping approach was also used for *E. coli* chromosome assembly using samtools/bcftools (Li *et al.*, 2009; Li, 2011). ECONIH4 (Weingarten *et al.*, 2018) was used as a reference genome for all *E. coli* isolates in this study. The plasmid gene synteny map was made using an in-house pipeline. The annotated sequence of the reference pOXA-48 plasmid (GenBank record JN626286, identical to RefSeq record NC_019154) (Poirel, Bonnin and Nordmann, 2012) was downloaded from GenBank (*GenBank Overview*, no date). A customised blast database containing all the samples was created and each gene annotated in the reference plasmid was then blasted (*BLAST: Basic Local Alignment Search Tool*, no date) against this database. Gene synteny was reconstructed using in-house programs and plotted using the genoplotR package (*BLAST: Basic Local Alignment Search Tool*, no date, *genoplotr*, no date) in R (*R: The R Project for Statistical Computing*, no date). A plasmid phylogenetic tree was built using the alignment resulting from reference based mapping using pOXA-48 as reference(Poirel, Bonnin and Nordmann, 2012). The tree was built using PhyML (Guindon *et al.*, 2010) in the analysis suite SeaView (Gouy, Guindon and Gascuel, 2010). We used GTR model (*Some Mathematical Questions in Biology*, no date; Yang, 1994; Zharkikh, 1994) with default PhyML options. Nucleotide variants in the plasmid sequence were identified using the seg.sites() command in the ape R package (‘Analyses of Phylogenetics and Evolution [R package ape version 5.3]’, no date) ClonalFrameML (Didelot and Wilson, 2015) with default options was used to infer the plasmid tree.

For the comparative analysis of ST399 *E. coli* isolates, all available sequences of *E. coli* ST399 were downloaded from EnteroBase (*Enterobase*, no date). These reads were processed following the same analysis pipeline as described for outbreak isolates.

### Expected number of mutations in p-OXA48 plasmid

The expected number of mutations is estimated from standard population genetics theory to be Nmut = μLΔt (Kimura, 1983). *E. coli* mutation rate is assumed to be 2.26·10^−7^ per site per year (Reeves *et al.*, 2011). pOXA-48 length is 61881 bp (Poirel, Bonnin and Nordmann, 2012). *E. coli* generation time being of the order of a few hours, we approximated 7 generations a day (Lenski *et al.*, 1991; Poirel, Bonnin and Nordmann, 2012). During the 15 years elapsed from the sampling of the reference pOXA-48 in 2001 and the sampling reported in this study in 2016, some 40,000 generations have passed (Lenski *et al.*, 1991).

Using these numbers the expected number of mutations is 0.209 (±0.44) or ≦2 mutations at 95% confidence interval. This estimate assumes the pOXA-48 reference as a direct ancestor to the present outbreak, and defines a lower limit to the number of expected mutations.

### Neutral model of inter-bacterial-host conjugation

Conjugation is modelled as a homogeneous Markov process between any two bacterial hosts. We assume: (i) that the taxonomic classification of the hosts is neutral, (ii) that the host populations have all similar abundances, (iii) that the epidemics started with a single host and (iv) the number of potential hosts is finite and equal to the ones in the samples at hand. In this study, we have considered each different ST within each observed species to be a different host. For this purpose, we consider equally different STs that belong to the same species or to different species. Therefore we have a total of 21 different potential hosts. The Markov process generator matrix M describing the instantaneous conjugation probability between different hosts in the Markov chain is therefore a 21 × 21 isotropic matrix. The host neutrality assumption (i) implies that the matrix depends on a single parameter, the conjugation rate *r*. More specifically, including the requirement that each row sums to 0, *M* has values − *r* on the diagonal elements and *r*/20 off-diagonal. The assumption of host neutrality combined with the assumption that the epidemics started with a single host implies that the most recent common ancestor (MRCA) of the outbreak was a pOXA-48 in *E. coli* ST399 strain. Therefore, the initial state distribution of the Markov process was a vector with 1 in the element *E. coli* ST399 and 0 in all the other elements corresponding to the other hosts.

The dynamics of a Markov process with a finite number of states is fully determined by its generator matrix. Given the initial distribution π(0) = π(*MRCA*) at *t*_*0*_, i.e. at the time of diversification of the MRCA, the distribution at any following time *t* is given by π(*t*) = π_0_*e*^*Mt*^. The probability that over all the time passed since the most recent common ancestor the plasmid never conjugated out of *E. coli* ST399 is *e*^−*rt*^ while the probability that it conjugated out is 1 − *e*^−*rt*^.

The probability that the plasmid is found in *E. coli* ST399 at a time *t* after *t*_*0*_ is the projection on *E. coli* ST399 of the exponential of matrix M:

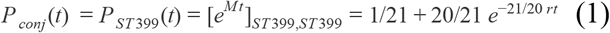

as π_*ST* 399_(0) = 1. The first term is time independent and corresponds to the stationary asymptotic state of the system, while the second term is time dependent, decreasing with time from *t*_*0*_.

The frequentist probability that the plasmid conjugates out of *E. coli* ST399 can be computed in our sample: *E. coli* ST399 is not the host in 20 of the 55 samples in this study, therefore this probability is 0.36. Equating this number to *e*^−*rt*^, we get *rt* = *ln* 0.36. Using this point estimate, we are able to estimate the probability that the plasmid is found in *E. coli* ST399 as a result of it conjugating back to it from another bacterial host in which it has previously conjugated during the time elapsed from the MRCA. We compute this probability as the probability of finding pOXA-48 in *E. coli* ST399 (Equation 1) minus the probability that it never conjugated out *e*^−*rT*^. It is ≃ 0.033. So there is around 3% of chance that we find p-OXA48 in a *E.coli* ST399 as a result of its conjugation from another bacterial host, and 97% probability that it comes either from vertical inheritance or conjugation from another *E.coli* ST399.

### Estimation of pOXA-48 conjugation rate

The evolution model described in “Neutral model of inter-bacterial-host conjugation” can be seen as a special case of a Wright-Fisher model (Hein, Schierup and Wiuf, 2004; Wakeley, 2009). This mathematical parallel allows us to apply a coalescent approach to determine the conjugation rate in our sample. The coalescent approach consists in applying a Kingman coalescent (Kingman, 1982) with conjugation instead of mutations happening at a constant rate along the tree branches. Conjugation changes the host from *E. coli* ST399, which we have seen is the initial host of the epidemics, to any other host. We make use of the infinite sites model approximation as it does not clash with the assumption we made of having a finite number of hosts. The non-ST399 *E. coli* hosts in our sample are more than 1/3, and, more importantly, they are all different from one another. This gives an effective decreasing probability that the plasmid conjugates twice in the same non-ST399 *E. coli* host(p<1/55~0.01), which justifies using the infinite sites model approximation (Kingman, 1982; Wakeley, 2009).

The probability of a conjugation event per unit time is *r*_*conj*_*dt*. Integrating over the tree height, the conjugation probability is 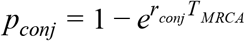. On the other end *p*_*conj*_ computed in a frequentist manner from our sample is 20/55 = 0.36

A dated phylogeny of the *E. coli* in the sample was obtained using BactDating (Didelot *et al.*, 2018). It was used to infer the the time since the MRCA (TMRCA) of the *E. coli* ST399 in the sample using a mutation rate per site per year of 2.26 10^−7^ (Reeves *et al.*, 2011). The estimation puts TMRCA around September 2014, so around 2 years before the onset of this outbreak. Using this point estimate, we infer a conjugation rate of *r*_*conj*_ ≃ 0.23 (±0.09) per lineage per year.

## Supporting information

Supplementary Table with Info about the samples

## Acknowledgements

A.L, X.D. and J.P thank the UK National Institute for Health Research Health Protection Research Unit (NIHR HPRU) in Modelling Methodology at Imperial College London in partnership with Public Health England (PHE) for funding (grant HPRU-2012–10080).

---

**Supplementary Figure 1:**
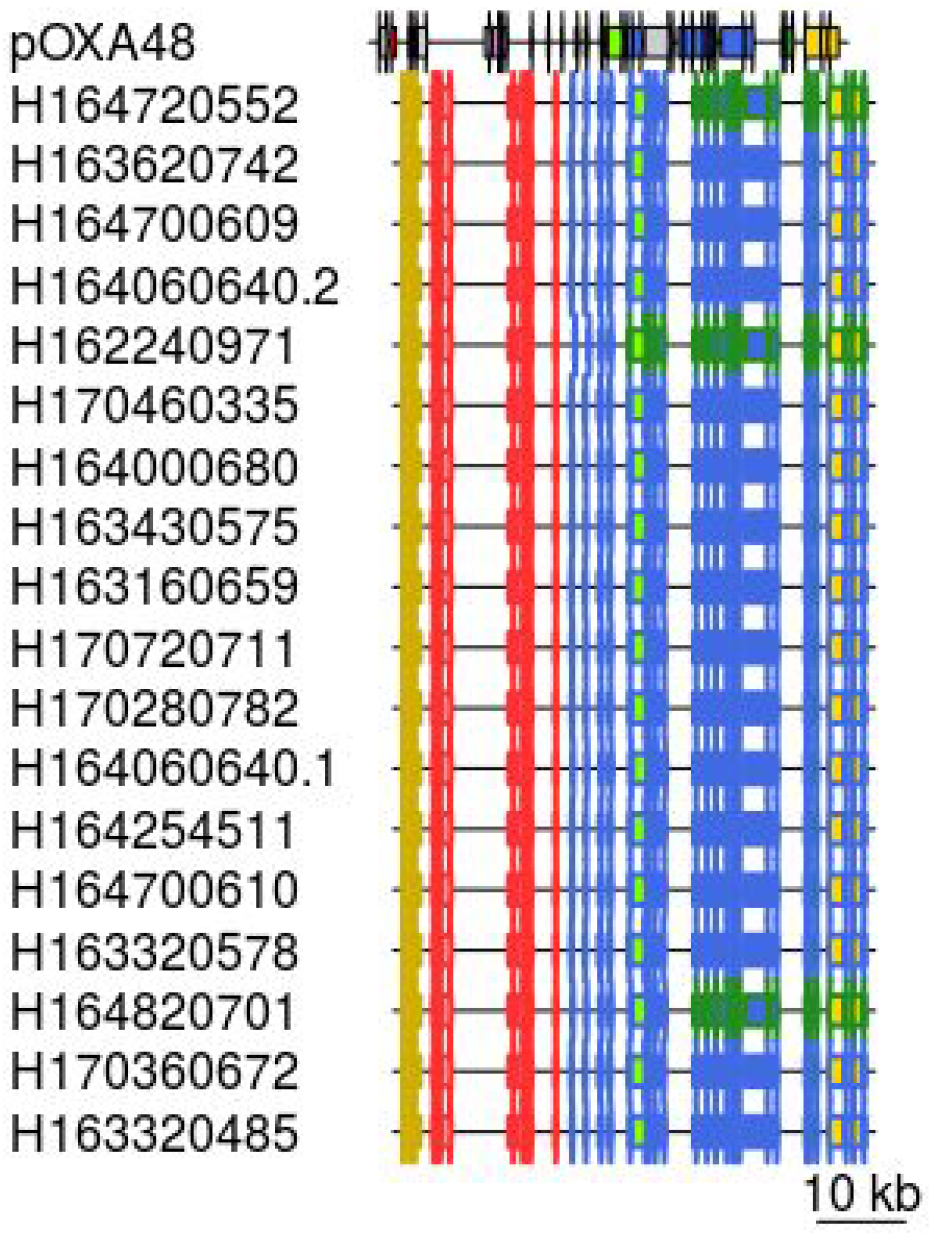
Plasmids gene content. The border color represents the contig (we have either 2 or 3 colors so it means that we find all the plasmid in either 2 or 3 contigs). The filling color is the same as in the reference plasmid (figure 3 a) and distinguishes different genes. In red OXA48. In purple the par genes (parA and parB), in light green the mob genes (mobA, mobB, mobC), in blue the tra genes, in dark green the reb genes (repA, repB, repC), in yellow the trb genes (trbA, trbB, trbC genes).

**Supplementary Figure 2.**
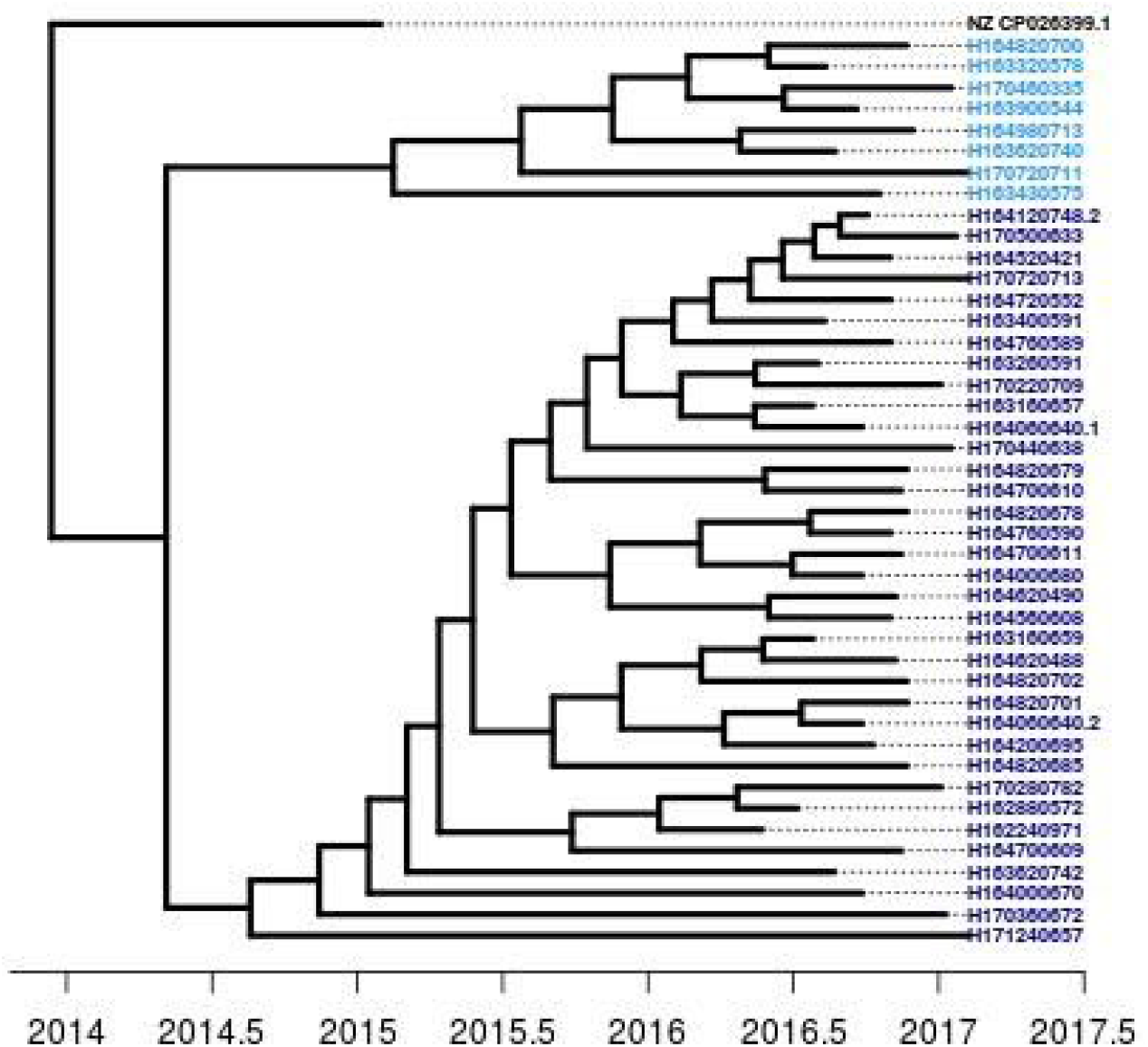
Dated phylogeny of all E. coli samples obtained with Bactdating. Most recent common ancestor dates to December 2013.

## References

‘Analyses of Phylogenetics and Evolution [R package ape version 5.3]’ (no date). Comprehensive R Archive Network (CRAN). Available at: https://CRAN.R-project.org/package=ape (Accessed: 26 May 2020).

Bankevich, A. et al. (2012) ‘SPAdes: a new genome assembly algorithm and its applications to single-cell sequencing’, Journal of computational biology: a journal of computational molecular cell biology, 19(5), pp. 455–477.

BLAST: Basic Local Alignment Search Tool (no date). Available at: https://blast.ncbi.nlm.nih.gov/Blast.cgi (Accessed: 26 May 2020).

Bonnin, R. A. et al. (2013) ‘Comparative genomics of IncL/M-type plasmids: evolution by acquisition of resistance genes and insertion sequences’, Antimicrobial agents and chemotherapy, 57(1), pp. 674–676.

Center for Genomic Epidemiology (no date a). Available at: http://www.genomicepidemiology.org/ (Accessed: 26 May 2020).

Center for Genomic Epidemiology (no date b). Available at: http://www.genomicepidemiology.org/ (Accessed: 26 May 2020).

Clausen, P. T. L. C., Aarestrup, F. M. and Lund, O. (2018) ‘Rapid and precise alignment of raw reads against redundant databases with KMA’, BMC bioinformatics, 19(1), p. 307.

Clermont, O. et al. (2013) ‘The Clermont Escherichia coli phylo-typing method revisited: improvement of specificity and detection of new phylo-groups’, Environmental microbiology reports, 5(1), pp. 58–65.

Dale, A. P. and Woodford, N. (2015) ‘Extra-intestinal pathogenic Escherichia coli (ExPEC): Disease, carriage and clones’, The Journal of infection, 71(6), pp. 615–626.

David, S. et al. (no date) ‘Genomic analysis of carbapenemase-encoding plasmids from Klebsiella pneumoniae across Europe highlights three major patterns of dissemination’. doi: 10.1101/2019.12.19.873935.

Didelot, X. et al. (2018) ‘Bayesian inference of ancestral dates on bacterial phylogenetic trees’, Nucleic acids research, 46(22), p. e134.

Didelot, X. and Wilson, D. J. (2015) ‘ClonalFrameML: efficient inference of recombination in whole bacterial genomes’, PLoS computational biology, 11(2), p. e1004041.

Enterobase (no date). Available at: https://enterobase.warwick.ac.uk/ (Accessed: 26 May 2020).

Ferretti, L. et al. (2017) ‘Decomposing the Site Frequency Spectrum: The Impact of Tree Topology on Neutrality Tests’, Genetics, 207(1), pp. 229–240.

Findlay, J., Hopkins, K. L., Alvarez-Buylla, A., et al. (2017) ‘Characterization of carbapenemase-producing Enterobacteriaceae in the West Midlands region of England: 2007-14’, Journal of Antimicrobial Chemotherapy, p. dkw560. doi: 10.1093/jac/dkw560.

Findlay, J., Hopkins, K. L., Loy, R., et al. (2017) ‘OXA-48-like carbapenemases in the UK: an analysis of isolates and cases from 2007 to 2014’, Journal of Antimicrobial Chemotherapy, pp. 1340–1349. doi: 10.1093/jac/dkx012.

Frazão, N. et al. (2019) ‘Horizontal gene transfer overrides mutation in Escherichia coli colonizing the mammalian gut’, Proceedings of the National Academy of Sciences, pp. 17906–17915. doi: 10.1073/pnas.1906958116.

GenBank Overview (no date). Available at: https://www.ncbi.nlm.nih.gov/genbank/ (Accessed: 26 May 2020).

genoplotr (no date). Available at: http://genoplotr.r-forge.r-project.org/ (Accessed: 26 May 2020).

Gouy, M., Guindon, S. and Gascuel, O. (2010) ‘SeaView version 4: A multiplatform graphical user interface for sequence alignment and phylogenetic tree building’, Molecular biology and evolution, 27(2), pp. 221–224.

Guindon, S. et al. (2010) ‘New algorithms and methods to estimate maximum-likelihood phylogenies: assessing the performance of PhyML 3.0’, Systematic biology, 59(3), pp. 307–321.

Hasman, H. et al. (2014) ‘Rapid whole-genome sequencing for detection and characterization of microorganisms directly from clinical samples’, Journal of clinical microbiology, 52(1), pp. 139–146.

Hein, J., Schierup, M. and Wiuf, C. (2004) Gene Genealogies, Variation and Evolution: A primer in coalescent theory. Oxford University Press, USA.

Hernández-García, M. et al. (2019) ‘Intestinal co-colonization with different carbapenemase-producing isolates is not a rare event in an OXA-48 endemic area’, EClinicalMedicine, 15, pp. 72–79.

Kimura, M. (1983) ‘The Neutral Theory of Molecular Evolution’. doi: 10.1017/cbo9780511623486.

Kingman, J. F. C. (1982) ‘The coalescent’, Stochastic Processes and their Applications, pp. 235–248. doi: 10.1016/0304-4149(82)90011-4.

Kurtz, S. et al. (2004) ‘Versatile and open software for comparing large genomes’, Genome biology, 5(2), p. R12.

Lanza, V. F. et al. (2014) ‘Plasmid Flux in Escherichia coli ST131 Sublineages, Analyzed by Plasmid Constellation Network (PLACNET), a New Method for Plasmid Reconstruction from Whole Genome Sequences’, PLoS Genetics, p. e1004766. doi: 10.1371/journal.pgen.1004766.

Larsen, M. V. et al. (2014) ‘Benchmarking of Methods for Genomic Taxonomy’, Journal of Clinical Microbiology, pp. 1529–1539. doi: 10.1128/jcm.02981-13.

Lenski, R. E. et al. (1991) ‘Long-Term Experimental Evolution in Escherichia coli. I. Adaptation and Divergence During 2,000 Generations’, The American Naturalist, pp. 1315–1341. doi: 10.1086/285289.

León-Sampedro, R. et al. (no date) ‘Dissemination routes of the carbapenem resistance plasmid pOXA-48 in a hospital setting’. doi: 10.1101/2020.04.20.050476.

Li, H. et al. (2009) ‘The Sequence Alignment/Map format and SAMtools’, Bioinformatics, 25(16), pp. 2078–2079.

Li, H. (2011) ‘A statistical framework for SNP calling, mutation discovery, association mapping and population genetical parameter estimation from sequencing data’, Bioinformatics, 27(21), pp. 2987–2993.

MacLean, R. C. and San Millan, A. (2019) ‘The evolution of antibiotic resistance’, Science, 365(6458), pp. 1082–1083.

Mahon, B. M. et al. (2019) ‘Detection of OXA-48-like-producing Enterobacterales in Irish recreational water’, Science of The Total Environment, pp. 1–6. doi: 10.1016/j.scitotenv.2019.06.480.

Mathers, A. J. et al. (2015) ‘Klebsiella pneumoniae Carbapenemase (KPC)-Producing K. pneumoniae at a Single Institution: Insights into Endemicity from Whole-Genome Sequencing’, Antimicrobial Agents and Chemotherapy, pp. 1656–1663. doi: 10.1128/aac.04292-14.

Nordmann, P., Dortet, L. and Poirel, L. (2012) ‘Carbapenem resistance in Enterobacteriaceae: here is the storm!’, Trends in molecular medicine, 18(5), pp. 263–272.

Norman, A., Hansen, L. H. and Sørensen, S. J. (2009) ‘Conjugative plasmids: vessels of the communal gene pool’, Philosophical transactions of the Royal Society of London. Series B, Biological sciences, 364(1527), pp. 2275–2289.

Poirel, L. et al. (2011) ‘OXA-163, an OXA-48-Related Class D β-Lactamase with Extended Activity Toward Expanded-Spectrum Cephalosporins’, Antimicrobial Agents and Chemotherapy, pp. 2546–2551. doi: 10.1128/aac.00022-11.

Poirel, L., Bonnin, R. A. and Nordmann, P. (2012) ‘Genetic features of the widespread plasmid coding for the carbapenemase OXA-48’, Antimicrobial agents and chemotherapy, 56(1), pp. 559–562.

Poirel, L., Potron, A. and Nordmann, P. (2012) ‘OXA-48-like carbapenemases: the phantom menace’, The Journal of antimicrobial chemotherapy, 67(7), pp. 1597–1606.

Potron, A. et al. (2011) ‘Characterization of OXA-181, a Carbapenem-Hydrolyzing Class D β-Lactamase from Klebsiella pneumoniae’, Antimicrobial Agents and Chemotherapy, pp. 4896–4899. doi: 10.1128/aac.00481-11.

Pulss, S. et al. (2018) ‘Multispecies and Clonal Dissemination of OXA-48 Carbapenemase in Enterobacteriaceae From Companion Animals in Germany, 2009—2016’, Frontiers in Microbiology. doi: 10.3389/fmicb.2018.01265.

Rahmani, A.-M. et al. (2011) ‘LastZ: An Ultra Optimized 3D Networks-on-Chip Architecture’, 2011 14th Euromicro Conference on Digital System Design. doi: 10.1109/dsd.2011.26.

Reeves, P. R. et al. (2011) ‘Rates of mutation and host transmission for an Escherichia coli clone over 3 years’, PloS one, 6(10), p. e26907.

R: The R Project for Statistical Computing (no date). Available at: https://www.r-project.org/ (Accessed: 26 May 2020).

Sheppard, A. E. et al. (2016) ‘Nested Russian Doll-Like Genetic Mobility Drives Rapid Dissemination of the Carbapenem Resistance Gene blaKPC’, Antimicrobial agents and chemotherapy, 60(6), pp. 3767–3778.

Skalova, A. et al. (2017) ‘Molecular Characterization of OXA-48-Like-Producing Enterobacteriaceae in the Czech Republic and Evidence for Horizontal Transfer of pOXA-48-Like Plasmids’, Antimicrobial agents and chemotherapy, 61(2). doi: 10.1128/AAC.01889-16.

Some Mathematical Questions in Biology (no date) *Google Books*. Available at: https://books.google.com/books/about/Some_Mathematical_Questions_in_Biology.html?id=8aI1phhOKhgC (Accessed: 26 May 2020).

Stoesser, N. et al. (2013) ‘Predicting antimicrobial susceptibilities for Escherichia coli and Klebsiella pneumoniae isolates using whole genomic sequence data’, The Journal of antimicrobial chemotherapy. Oxford Academic, 68(10), pp. 2234–2244.

Turton, J. F. et al. (2016) ‘Clonal expansion of Escherichia coli ST38 carrying a chromosomally integrated OXA-48 carbapenemase gene’, Journal of medical microbiology, 65(6), pp. 538–546.

Wakeley, J. (2009) Coalescent Theory: An Introduction. Roberts Publishers.

Website (no date). Available at: https://github.com/tseemann/nullarbor (Accessed: 26 May 2020).

Weingarten, R. A. et al. (2018) ‘Genomic Analysis of Hospital Plumbing Reveals Diverse Reservoir of Bacterial Plasmids Conferring Carbapenem Resistance’, mBio, 9(1). doi: 10.1128/mBio.02011-17.

Wen-Hsiung, F. Y.-X. L. (1993) ‘Statistical tests of neutrality of mutations’, Genetics, 133(3), pp. 693–709.

Yang, Z. (1994) ‘Estimating the pattern of nucleotide substitution’, Journal of molecular evolution, 39(1), pp. 105–111.

Zharkikh, A. (1994) ‘Estimation of evolutionary distances between nucleotide sequences’, Journal of Molecular Evolution, pp. 315–329. doi: 10.1007/bf00160155.

